# Unconventional centromere architectures in *Tapirus indicus* reveal hotspots for satellite-free centromere formation in Perissodactyla

**DOI:** 10.1101/2025.10.09.681474

**Authors:** Marialaura Biundo, Francesca M. Piras, Edoardo Rapisarda, Oliver A. Ryder, Solomon G. Nergadze, Elena Giulotto, Eleonora Cappelletti

## Abstract

Centromeres, the chromosomal loci responsible for proper segregation during cell division, play a key role in genome evolution and speciation. While centromere function is highly conserved and epigenetically defined by CENP-A, the underlying DNA sequences are among the most rapidly evolving. Although mammalian centromeres are typically associated with satellite DNA, we previously showed that equids carry numerous satellite-free centromeres.

In this study, we investigated centromere and karyotype evolution in the endangered *Tapirus indicus*, a non-equid Perissodactyl with exceptional karyotypic plasticity. Through CENP-A ChIP-seq analysis on the same individual for which a near-gapless diploid genome assembly was available, we identified both canonical satellite-based centromeres and three satellite-free centromeres, emerging from centromere repositioning and representing the first such centromeres described in a non-equid Perissodactyl species. Comparative genomic analysis uncovered evolutionary hotspots for satellite-free centromere formation across Perissodactyla. Finally, analysis of CENP-B binding showed that *T. indicus* displays uncoupling between CENP-A and CENP-B, a feature previously observed only in equids. These findings reveal that high centromere plasticity is not unique to equids and support a broader model in which centromere plasticity and CENP-B uncoupling contribute to karyotype evolution in mammals.

## INTRODUCTION

Centromeres, the chromosomal loci that ensure proper segregation during cell division, play a crucial role in genome evolution and speciation, as well as in cancer and chromosome abnormalities. The centromere function is highly conserved across species and is epigenetically specified by the histone H3 variant CENP-A (Henikoff et al. 2001). Despite this functional conservation, the underlying DNA sequences are remarkably divergent and represent some of the most rapidly evolving regions in eukaryotic genomes (Plohl et al. 2008). In mammals, centromeric DNA is typically composed of highly repetitive satellite sequences, which has long posed a significant obstacle to the study of these chromosomal regions, leaving them among the most obscure parts of their genomes. Recent methods for assembling highly accurate and nearly complete genomes have begun to overcome these barriers, opening access to repetitive regions and enabling large-scale sequencing of neglected species crucial for evolutionary biology and biodiversity conservation (Rhie et al. 2021; Altemose et al. 2022; Gupta 2022; Nurk et al. 2022; Miga and Eichler 2023).

The remarkable variation in centromere organization among mammals is further exemplified by our discovery that equid species possess several functional centromeres completely devoid of satellite DNA (Wade et al. 2009; Piras et al. 2010; Nergadze et al. 2018; Cappelletti et al. 2022; Piras et al. 2022; Piras et al. 2023; Cappelletti et al. 2025a). Although a few satellite-free centromeres have been identified in other mammals in subsequent studies (Tolomeo et al. 2017; Hartley et al. 2025), the genus *Equus* remains unique in this respect, with species such as donkeys and zebras exhibiting satellite-free centromeres at more than half of their chromosomes.

This unusual centromere architecture appears to be a consequence of the recent and rapid evolution of the genus, which occurred within the last four million years and involved extensive karyotypic reshuffling. Two key mechanisms contributed to this reorganization and the formation of satellite-free centromeres: centromere repositioning - the shift of centromere position without accompanying chromosomal rearrangements - and chromosomal fusions, particularly centric ones (Carbone et al. 2006; Piras et al. 2009; Piras et al. 2010; Nergadze et al. 2018; Cappelletti et al. 2022; Piras et al. 2023). We recently suggested that the exceptional plasticity of equid centromeres may be linked to the evolutionary dissociation of CENP-A from CENP-B (Cappelletti et al. 2025b), the only known centromeric protein with sequence-specific DNA binding (Masumoto et al. 1989), which is implicated in centromere stability in humans and mice (Gamba and Fachinetti 2020).

The species of the genus *Equus* are the only extant members of the family Equidae, which belongs to the suborder Hippomorpha of the order Perissodactyla. The other living families of this order, Rhinocerotidae (rhinoceroses) and Tapiridae (tapirs), together form the suborder Ceratomorpha, which diverged from Hippomorpha approximately 56 million years ago. While equid karyotypes underwent rapid evolution, the karyotypes of tapirs and rhinoceroses have remained quite stable and closely resemble the hypothetical ancestral karyotype (Trifonov et al. 2008; Trifonov et al. 2012). The only exception is the Malayan tapir (*Tapirus indicus*), whose karyotype was restructured through a series of fusions, making it a unique case within Ceratomorpha (Trifonov et al. 2008; Trifonov et al. 2012). This endangered species is the only extant Asian tapir, whereas the other *Tapirus* species (*T. terrestris*, *T. bairdii*, and *T. pinchaque*) are native to South America (Houck et al. 2000).

In this work, we investigated centromere and karyotype evolution in *T. indicus*, demonstrating that satellite-free centromeres are present and coexist with canonical satellite-based centromeres. Both types were thoroughly characterized, taking advantage of the release of a high-quality diploid genome assembly that includes satellite sequences. We showed that the satellite-free centromeres originated through centromere repositioning events and identified hotspots for neocentromere formation across Perissodactyla. Furthermore, by analyzing the binding pattern of CENP-B, we identified *T. indicus* as a new case of uncoupling between CENP-A and CENP-B, further supporting a link between centromere reshuffling and the loss of CENP-B association. Altogether, this work provides new insights into the complexity and evolutionary dynamics of centromere organization within the rich diversity of mammalian genomes.

## RESULTS

### Chromosome number assignment to the *Tapirus indicus* genome assembly and search of satellite DNA families

A chromosome-level, haplotype-phased genome assembly of *Tapirus indicus* was recently produced by the Vertebrate Genome Project using PacBio HiFi and Dovetail Omni-C data from a female donor. The diploid assembly includes two haplotypes: mTapInd1.hap2, which serves as the reference assembly in NCBI, and mTapInd1.hap1. The original chromosomal numbering in this assembly is arbitrary and does not reflect the cytogenetically defined karyotype of *Tapirus indicus* (Houck et al. 2000; Trifonov et al. 2008; Trifonov et al. 2012). Since the chromosomal orthologies between *Tapirus indicus* and *Equus caballus* are well described (Trifonov et al. 2008), we assigned correct chromosome numbers to the mTapInd1.hap2 and mTapInd1.hap1 sequences by performing a whole-genome alignment with the T2T_TB horse assembly (Supplementary Fig. 1, Supplementary Table 1).

The nearly error-free and gapless nature of this assembly makes it an excellent resource for the detailed characterization of satellite DNA and centromere organization in this species. Since satellite families in *Tapirus indicus* had not been characterized, we used TAREAN (Novák et al. 2017) to *de novo* identify satellite repeats. As this pipeline relies on unassembled short reads, we generated a short-read whole-genome sequencing dataset (SRR35264171) from the same individual used for the diploid genome assembly. This analysis revealed that two satellite families, that we named TINSat1 and TINSat2, are present (Supplementary Table 2). TINSat1, the most abundant satellite family, has a 646 bp long consensus sequence with a GC content of 48.1%. In contrast, the consensus of TINSat2 is 2,688 bp long, has a higher GC content of 66.26% and is poorly represented in the analyzed reads. Genome-wide mapping of these satellite families in the diploid assembly (Fig. 1A, Supplementary Tables 3 and 4) confirmed that TINSat1 is the most abundant satellite repeat, spanning approximately 41 Mb in mTapInd1.hap1 (1.69% of the genome) and 32 Mb in mTapInd1.hap2 (1.32 % of the genome). TINSat2 covers 5 Mb in mTapInd1.hap1 and 5.5 Mb in mTapInd1.hap2 and accounts for 0.20% and 0.23% of the genome, respectively.

**Figure 1.**
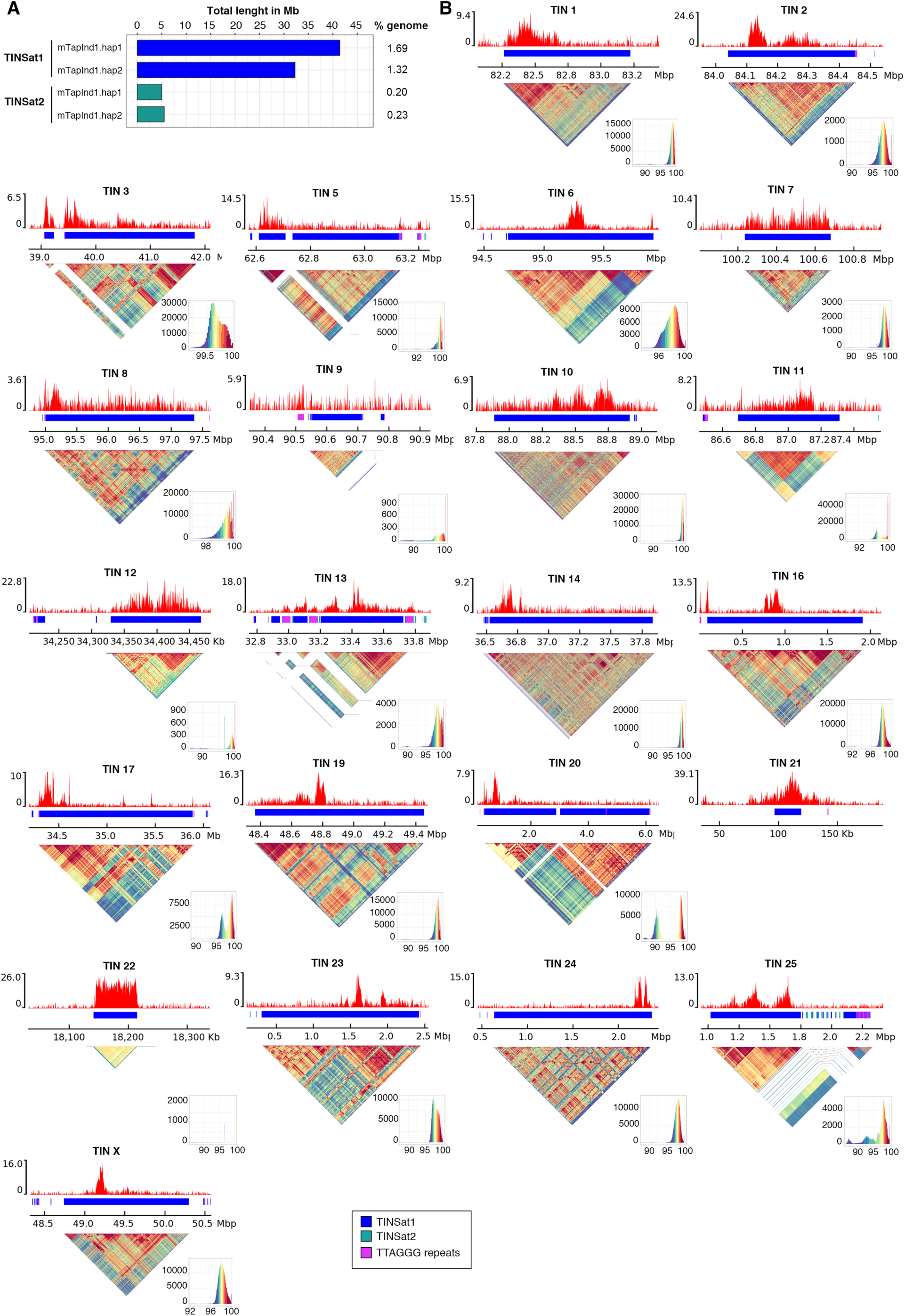
Satellite DNA families and satellite-based centromeres in the *Tapirus indicus* genome assembly. A) Barplots showing the total lengths (in megabases) of each satellite DNA family in haplotype 1 and haplotype 2. The genome percentage of each family is indicated on the right. B) Satellite-based centromeres of *T. indicus* in the haplotype 2 reference. For each centromere, the ChIP-seq profile of CENP-A is shown at the top. The y-axis reports the normalized read counts whereas the x-axis reports the coordinates on the mTapiInd1.hap2 assembly. The colored bar represents the satellite DNA array, with colors denoting different satellite families as indicated in the legend shown in the last page. The sequence identity maps obtained with ModDotPlot are reported at the bottom.

### Satellite-based and satellite-free centromeres in *Tapirus indicus*

To localize functional centromeres, we performed a ChIP-seq experiment on fibroblasts from the same individual that provided DNA for the genome assembly, using an anti-horse-CENP-A antibody previously developed by our group (Cappelletti et al. 2019) and effectively recognizing tapir CENP-A (Supplementary Fig. 2). By combining ChIP-seq data with the high-quality diploid genome assembly from the same donor, we minimized artifacts due to inter-individual variation and satellite array heterogeneity and were able to map the functional centromeres in all chromosomes. As shown in Figure 1B and Supplementary Fig. 3, the CENP-A binding domain is embedded within a TINSat1 array in 23 out 26 chromosomes, with array sizes ranging from 22 kb (TIN21) to 7.5 Mb (TIN14) and typically falling between 1 and 3 Mb. At most loci, CENP-A enrichment peaks cover only a few hundred kilobases of the megabase-sized arrays, indicating that the remaining portion of the arrays is pericentromeric. On chromosomes with satellite arrays shorter than 100 kb (TIN19 in haplotype 1, TIN22 in haplotype 2 and TIN21 in both haplotypes), the CENP-A enrichment peak spans the entire array. Interestingly, the CENP-A binding domains of both TIN21 homologs (Figure 1B and Supplementary Figure 3) extend beyond the 22 kb TINSat1 array, suggesting that the satellite may be in an early stage of expansion and that this centromere may be not yet fully “mature”. The shape of the enrichment peaks also varies considerably, ranging from Gaussian-like peaks (e.g. TIN6 in haplotype 2) to irregular profiles (e.g. TIN10 in haplotype 2). In some cases, such as TIN3 and TIN13 in both haplotypes, CENP-A enrichment appears diffuse across the array, possibly reflecting mapping bias in regions where the satellite arrays are highly homogeneous.

Neither satellite arrays nor ChIP-seq enrichment peaks were detected on chromosome 20 of haplotype 1, likely due to a local assembly gap. Supporting this hypothesis, haplotype 2 is ∼7.5 Mb longer than haplotype 1 for this chromosome, with additional sequence encompassing the first 6.2 Mb and a terminal segment of ∼1.5 Mb (Supplementary Fig. 4A). Both these regions in haplotype 2 harbor a CENP-A enrichment peak on satellite arrays and share sequence identity, even in portions not occupied by satellite DNA, suggesting that the missing sequence in haplotype 1 may have been partially misplaced at the end of haplotype 2 (Supplementary Fig. 4B).

To further explore the structure of the regions underlying CENP-A enrichment peaks, we used ModDotPlot - a heatmap-based visualization tool that evaluates tandem repeat architecture, sequence identity, and divergence (Sweeten et al. 2024). This analysis revealed that CENP-A binding domains are typically localized in the most homogeneous regions of TINSat1 arrays and are flanked by more divergent repeat units, supporting a model in which sequence uniformity contributes to centromere function and positioning (Fig. 1B, Supplementary Fig. 3).

While short stretches of TINSat2 and telomeric-like repeats are confined to the divergent pericentromeric regions of several chromosomes, both TIN13 homologs display an unusual centromeric architecture. Specifically, large blocks of telomeric-like repeats (40 kb each) are interspersed within a 1 Mb TINSat1 array. These telomeric blocks lie within the CENP-A binding domain, suggesting that telomeric-like sequences may be integrated components of the functional centromere on this chromosome (Fig. 1B, Supplementary Fig. 3). Another peculiar pattern was observed on TIN9 in haplotype 2, where a 20 kb CENP-A enrichment peak was detected over a telomeric-like repeat stretch, while the adjacent 200 kb TINSat1 array showed no enrichment. However, the short length and limited CENP-A enrichment in this region likely reflect an incomplete assembly.

The availability of both haplotypes allowed us to compare the positions of the CENP-A enrichment peaks on homologous chromosomes (Fig. 2A, Supplementary Fig. 5). It is important to note that variability in the satellite arrays between homologs adds complexity to the precise comparison of CENP-A binding domains. Though satellite arrays are generally conserved between homologs, sequence identity varies across their length. In several chromosomes (e.g. TIN1, TIN11 and TIN17), the CENP-A binding domains are localized within the regions of highest sequence identity between homologs (Fig. 2A, Supplementary Fig. 5). In contrast, in other cases (e.g. TIN23, TIN24 and TIN25), the sequences underlying the CENP-A binding domains are less conserved and the centromeric domains occupy different positions on the two homologs (Fig. 2A, Supplementary Fig. 5). Overall, homologous chromosomes exhibit higher satellite sequence similarity than non-homologous ones, although inter-chromosomal similarity remains relatively high, particularly within specific chromosome groups (Fig. 2B).

**Figure 2.**
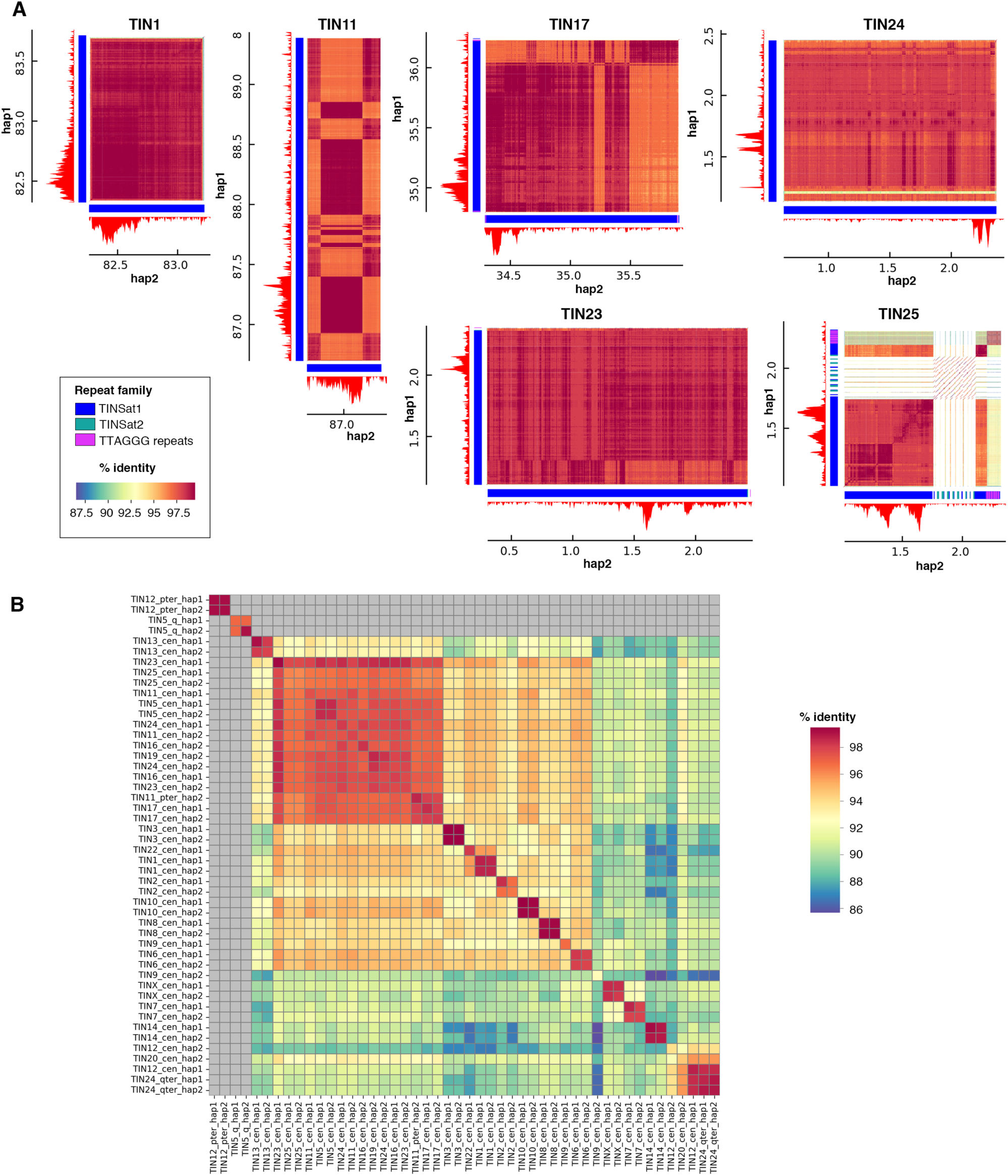
Variation in satellite DNA and CENP-A binding domains at satellite-based centromeres. A) Comparison of satellite DNA organization and CENP-A binding domains across homologous chromosomes. For each centromere, ChIP-seq profiles of CENP-A in mTapiInd1.hap1 (hap1) and mTapiInd1.hap2 (hap2) are shown. Colored bars beneath the CENP-A enrichment peaks represent satellite DNA arrays, with colors corresponding to different satellite families as defined in the legend. All array lengths are drawn to scale. Sequence identity maps between the two haplotypes, generated using ModDotPlot, are displayed in the center. The color scale for the identity heatmap is also provided in the legend on the first page. B) Heatmap showing the percentage of identity among satellite array regions between haplotypes. Each cell represents the average identity (%) calculated using ModDotPlot. Gray cells indicate comparisons with no detectable identity. The long non-centromeric loci (5q, 11pter, 12pter, 24qter) are included.

While most centromeres are associated with satellite DNA, ChIP-seq analysis revealed that chromosomes TIN4, TIN15, and TIN18 harbor satellite-free centromeres (Fig. 3A). For each centromere, CENP-A enrichment peaks were identified in both haplotypes in homologous regions (red peaks in Fig. 3A). We previously demonstrated that homologs often display CENP-A binding domains at different positions (Purgato et al. 2015; Nergadze et al. 2018; Cappelletti et al. 2023). However, due to the high sequence conservation between the two haplotypes, conventional mapping approaches were insufficient to distinguish the CENP-A binding domains of the two homologs (red peaks in Fig. 3A). To overcome this limitation, we developed a strategy based on Single Nucleotide Variants (SNVs) to map the ChIP-seq reads in a haplotype-specific manner. After identifying SNVs (Supplementary Table 5), we retained only reads overlapping these positions with no mismatches and used them to generate haplotype-specific enrichment profiles (orange peaks in Fig. 3A). This approach enabled clear distinction of the CENP-A binding domains on each homolog.

**Figure 3.**
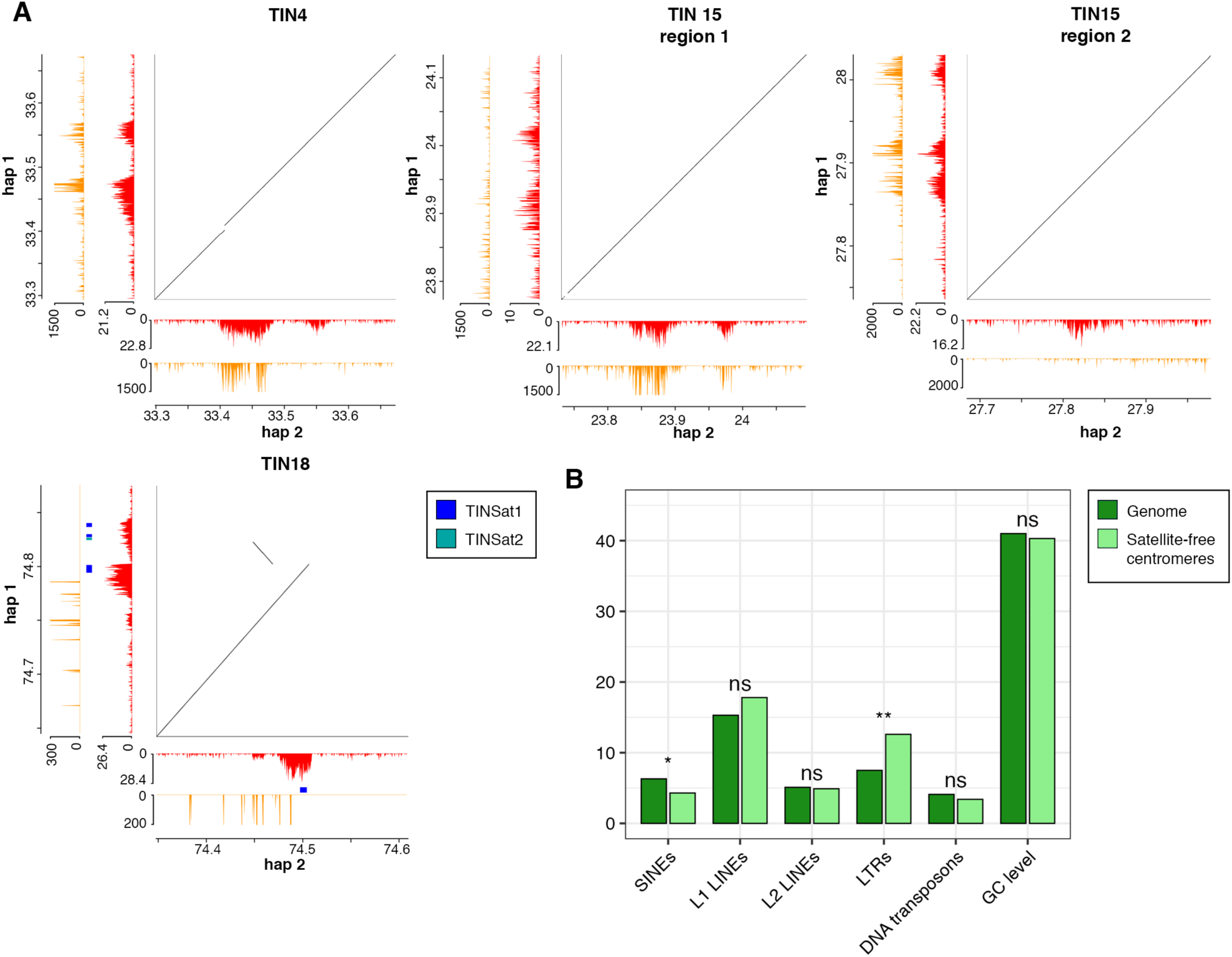
Satellite-free centromeres in *Tapirus indicus*. A) ChIP-seq profiles of CENP-A enrichment for each centromere in mTapiInd1.hap1 (hap1) and mTapiInd1.hap2 (hap2). Profiles obtained using conventional mapping are shown in red, while those generated with the SNV-based approach are shown in orange. Sequence identity dotplots between the two haplotypes are displayed in the center. Colored bars beneath the CENP-A peaks of TIN18 indicate short stretches of satellite DNA repeats, with colors corresponding to satellite families defined in the legend. B) Sequence composition of satellite-free centromeres. The genomic regions underlying the CENP-A enrichment peaks were analyzed for GC content, SINEs, L1 LINEs, L2 LINEs, LTRs, and DNA transposons. Values were compared to genome-wide averages. Differences in SINEs and LTRs are statistically significant (p-values 0.024 and 0.009, respectively). Differences in L1 LINEs, L2 LINEs, DNA transposons, and GC content were not significant (ns).

On TIN4, a main enrichment peak (90 kb) and a secondary peak (45 kb), separated by a 40 kb region lacking CENP-A binding were detected. SNV analysis demonstrated that the CENP-A binding domain of haplotype 1 spans part of the 90 kb region and includes the 45 kb secondary peak. The peak on haplotype 2 covers the entire 90 kb region but does not extend to the secondary peak.

In TIN15 there are two distinct CENP-A enriched regions located 3.7 Mb apart. These two regions are located on different homologs: region 1 maps to the haplotype 2 homolog and region 2 maps to the haplotype 1 homolog (Fig. 3A).

On TIN18, CENP-A enrichment was detected at corresponding positions on both homologs with peak domains spanning 100 kb on haplotype 1 and 70 kb on haplotype 2 (Fig. 3A). Interestingly, each domain is largely satellite-free, containing only a few scattered arrays of satellite repeats, each spanning just a few kilobases (Fig. 3A, Supplementary Tables 3 and 4). Given their negligible size relative to the overall CENP-A binding regions, the TIN18 centromere can be classified as satellite-free.

RepeatMasker analysis comparing the sequence composition of satellite-free centromeres to genome-wide averages revealed that the regions underlying CENP-A binding domains are enriched in LTR elements, while SINE elements are slightly under-represented (Fig. 3B). GC content, as well as the abundance of LINE1, LINE2, and DNA transposons, does not differ from genome-wide averages.

### Genome-wide landscape of satellite DNA and telomeric repeats in *Tapirus indicus*

A global overview of centromere and satellite organization across the *T. indicus* chromosomes is depicted in Figure 4, where the positions of centromeres and satellite arrays in both haplotypes are shown. The satellite-based centromeres are primarily composed of TINSat1, which makes up the majority of the (peri)centromeric satellite arrays.

**Figure 4.**
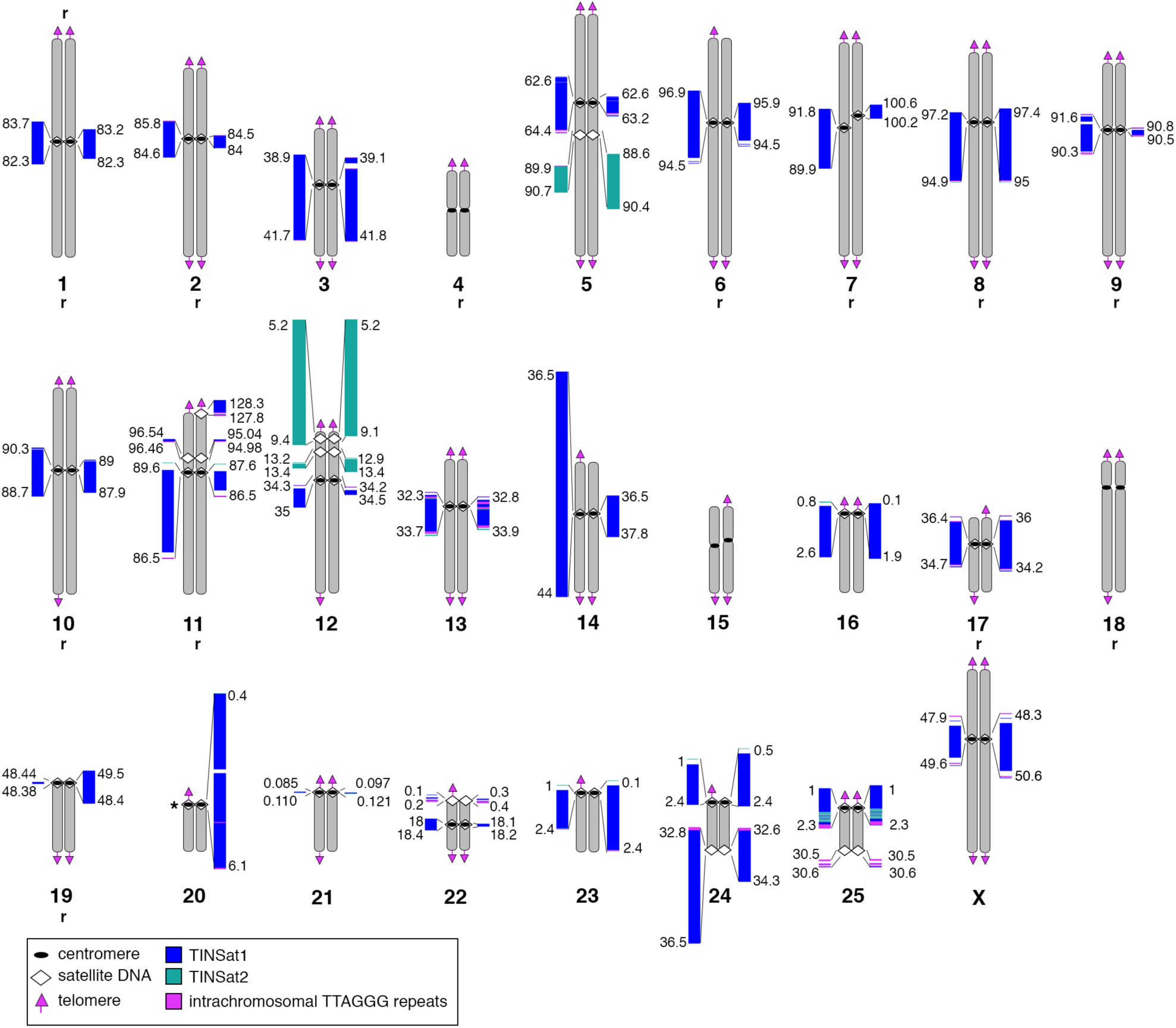
Satellite DNA and centromere annotation in the *Tapirus indicus* genome assembly. B) Schematic representation of homologous chromosome pairs, with centromeres (black ovals), satellite DNA arrays (white lozenges), and telomeres (pink arrows) indicated. For each pair, haplotype 1 is shown on the left and haplotype 2 on the right. A color-coded map of satellite DNA arrays is shown for each satellite locus, with array lengths drawn to scale. Only non-centromeric satellite arrays longer than 100 kb are included. Intrachromosomal TTAGGG repeats associated with satellite arrays are also shown. Chromosomes that are reversed in orientation in the assembly relative to their actual orientation are marked with an “r”. The asterisk indicates the centromere of the TIN20 chromosome, which is misassembled in haplotype 2 but identified as satellite-based according to cytogenetic analysis.

Extended non-centromeric satellite loci were identified at TIN5q, TIN12p, TIN22p and TIN25q. The loci on TIN5q and TIN12p correspond to the only large genomic arrays of the TINSat2 family, which is otherwise minimally represented at (peri)centromeric regions. The remaining non-centromeric satellite loci are shorter and composed primarily of TINSat1. The non-centromeric satellite array at the p terminus of TIN11 is present only in haplotype 2, suggesting structural variation between homologous chromosomes or mis-assembly of this region. In addition to the extended arrays shown in Fig. 4, short sequences homologous to satellite repeats were also found at several interstitial non-centromeric sites (Supplementary Tables 3 and 4).

The satellite DNA distribution identified in the diploid genome assembly was confirmed by FISH analysis on metaphase chromosomes using total genomic DNA as a probe (Supplementary Fig. 8), a technique we previously showed to be effective in detecting highly repetitive DNA due to its distinct hybridization kinetics compared to single-copy sequences (Piras et al. 2010; Piras et al. 2023). While most primary constrictions displayed hybridization signals, no signals were detected at the primary constrictions of chromosomes TIN4, TIN15 and TIN18 (Supplementary Fig. 8), consistent with their satellite-free status (Fig. 3A). TIN21 and TIN22 also lacked detectable signals, in line with the very short satellite arrays reported in the assembly (Fig. 4, Supplementary Tables 3 and 4), which likely fall below the FISH detection threshold. Non-centromeric signals were also observed at the p-terminus of TIN11 of one homolog, confirming the presence of satellite DNA on haplotype 2 only. In contrast, hybridization signals were observed on both TIN20 homologs, even though the satellite array and centromeric region are missing from haplotype 1 in the assembly, confirming the hypothesis of an assembly gap in this region.

To complete the sequence analysis, we searched for TTAGGG telomeric repeats in both haplotypes (Supplementary Tables 6 and 7). Telomeric arrays, typically spanning a few kilobases, were identified at most chromosomes (Fig. 4, Supplementary Tables 6 and 7). Longer telomeric-like arrays - ranging from a few kilobases to several tens of kilobases - were found either flanking or embedded within satellite DNA arrays, primarily within TINSat1 (Fig. 4, Supplementary Tables 6 and 7). FISH with a telomeric repeat oligonucleotide probe confirmed the presence of telomeric-like repetitions at several primary constrictions (Supplementary Fig. 9). Short interstitial telomeric sequences (ITSs), ranging from a few base pairs to approximately 1 kb, were scattered throughout the genome, as commonly observed in mammals (Nergadze et al. 2004; Nergadze et al. 2007; Ruiz-Herrera et al. 2008; Santagostino et al. 2020; Sola et al. 2021). These short ITSs were not detectable by FISH and are not reported in Figure 4.

### Hotspots for neocentromere formation in Perissodactyla

The three satellite-free centromeres identified in *T. indicus* represent the first such cases reported in Perissodactyla species other than equids. To explore their origin, we first performed whole-chromosome alignments between these chromosomes and the corresponding orthologs identified in the chromosomal-level assemblies of two rhinoceros (*Ceratotherium simum*, *Diceros bicornis*), two other tapirs (*Tapirus terrestris*, *Tapirus bairdii*), and four equids (*Equus caballus*, *Equus asinus*, *Equus burchelli*, *Equus grevyi*) (Supplementary Fig. 10). After the identification of orthologous chromosomes in the Perissodactyla species, it was important to localize the position of their centromeres. However, while in equids the position of centromeres was previously well defined (Wade et al. 2009; Piras et al. 2010; Nergadze et al. 2018; Cappelletti et al. 2022; Cappelletti et al. 2025b), for rhinoceroses and other tapirs, centromere positions have so far only been inferred from cytogenetic data (Houck et al. 2000; Trifonov et al. 2008; Trifonov et al. 2012) and satellite families remained uncharacterized. To complement these data, we used TAREAN to identify satellite sequences from publicly available whole-genome sequence datasets (Supplementary Table 8), and mapped them onto the orthologous chromosomes (Supplementary Tables 9-12). In the high-quality, long-read assemblies of *C. simum* and *D. bicornis*, megabase-sized satellite arrays, likely corresponding to the characteristic heterochromatic blocks of rhinoceros (peri)centromeres, were detected. In contrast, in the short-read assemblies of *T. terrestris* and *T. bairdii*, the satellite arrays were only partially assembled or entirely missing. We then integrated chromosomal alignments with centromere positions and satellite distribution to reconstruct the evolutionary history of these orthologous chromosome groups (Fig. 5A). A detailed description of the comparative analysis is reported in the Supplementary text. Briefly, we demonstrated that the satellite-free centromeres of TIN4, TIN15 and TIN18 emerged from centromere repositioning. A surprising finding was that TIN4 and its horse ortholog ECA11 (Wade et al. 2009) carry a satellite-free centromere at the same position (Fig. 5B). Similarly, the 4.2 Mb region encompassing the two distant TIN15 epialleles is orthologous to the region in the donkey genome where the centromere of donkey chromosome 8 (EAS8) (Nergadze et al. 2018) emerged following a centromere repositioning event (Fig. 5C). In the donkey, this centromere harbors DNA duplications that were expanded specifically in the donkey lineage and likely represent early stages of chromosome-specific centromeric satellite DNA formation (Nergadze et al. 2018). As described in detail in the Supplementary text, given the evolutionary distance between the Tapiridae and Equidae families (Fig. 5A), these results suggest that independent centromere repositioning events occurred in the *T. indicus* and *E. caballus* lineages to generate the satellite-free centromeres on TIN4 and ECA11. Similarly, independent centromere repositioning events occurred in the *T. indicus* and *E. asinus* lineages to generate the satellite-free centromeres on TIN15 and EAS8. The emergence of satellite-free centromeres at orthologous positions in different lineages indicates the presence of hotspots for neocentromere formation in the genomes of Perissodactyla.

**Figure 5.**
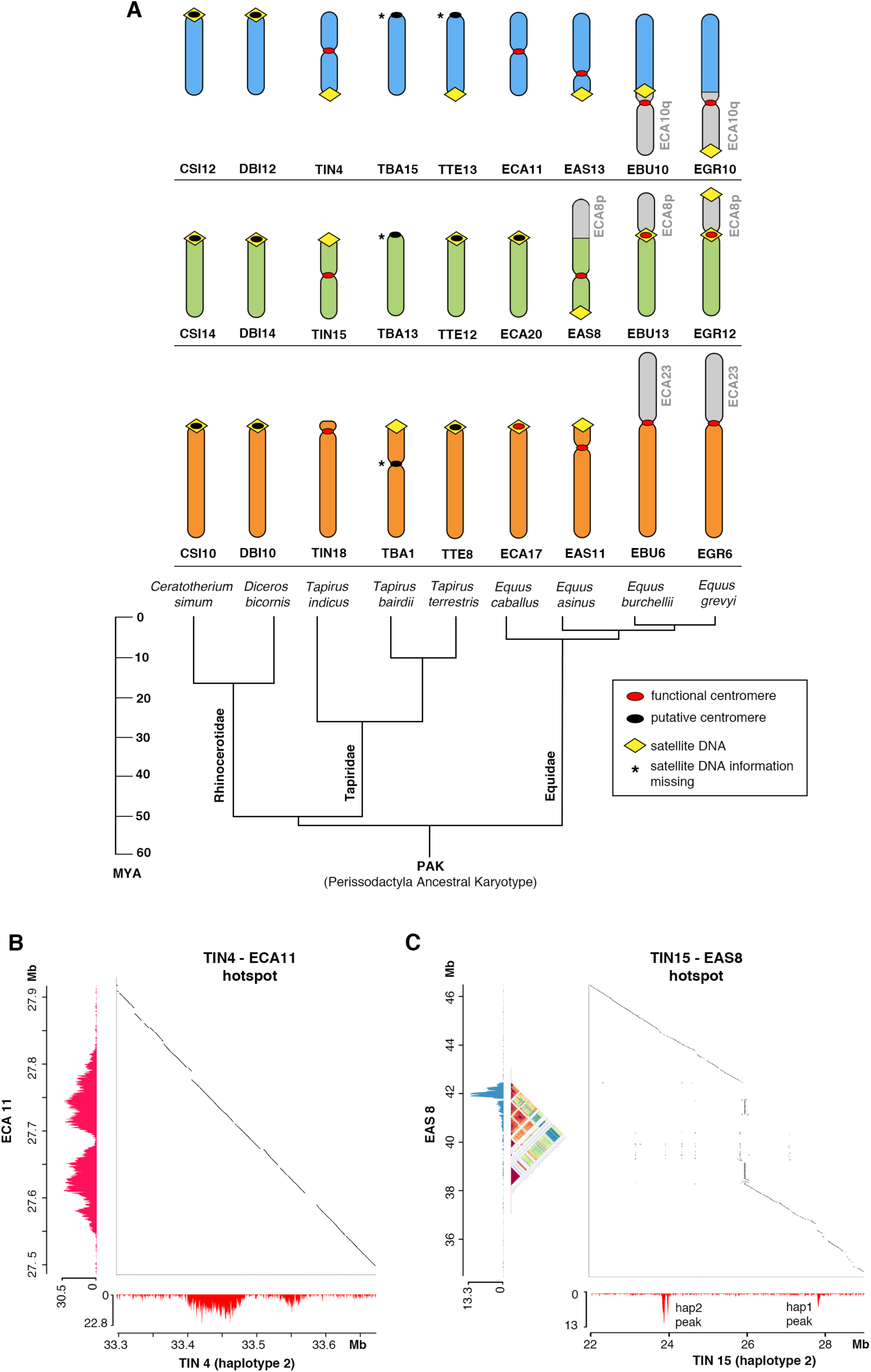
Hotspots for neocentromere formation in Perissodactyla. A) Comparison between tapir chromosomes carrying satellite-free centromeres and their orthologous chromosomes in other perissodactyl species. Colors indicate orthologous chromosomal segments. The positions of satellite arrays, including very short stretches, identified in the respective genome assemblies are shown as yellow lozenges. The positions of functional centromeres characterized at the molecular level (red ovals) are available for *T. indicus*, *E. caballus*, *E. asinus*, *E. grevyi* and *E. burchelli*. For *D. bicornis*, *C. simum*, *T. terrestris* and *T. bairdii*, the positions of putative centromeres inferred from karyotype analyses are shown (black ovals). Centromeres for which information on the presence of satellite DNA is unavailable are marked with asterisks. Below each set of orthologous chromosomes, the current phylogenetic tree of Perissodactyla is shown, based on Trifonov et al. (2008, 2012). B) Comparison of the satellite-free CENP-A binding domains in *T. indicus* chromosome 4 and *E. caballus* chromosome 11. ChIP-seq profiles of CENP-A for both species are shown, with sequence identity maps between the two orthologous regions displayed in the center. C) Comparison of the satellite-free CENP-A binding domains in *T. indicus* chromosome 15 and *E. asinus* chromosome 8. ChIP-seq profiles of CENP-A are shown for both species. Below the CENP-A peak in *E. asinus*, the identity plot generated with ModDotPlot highlights the region of DNA duplication. Sequence identity maps between the orthologous regions are displayed in the center. A detailed description of this figure is reported in the Supplementary text.

The third *T. indicus* satellite-free centromere, on TIN18, occurred in this lineage only.

### CENP-B binding sites in *Tapirus* species

CENP-B is the only centromeric protein known to bind a specific DNA motif (CENP-B box), which is present in major centromeric satellites of several mammals. However, we recently demonstrated that, in equids, CENP-B is uncoupled from centromere function (Cappelletti et al. 2025b).

To investigate CENP-B in *Tapirus indicus*, we first identified the CENP-B gene in the diploid genome and found that its protein coding sequence contains a conserved DNA-binding domain, suggesting that it is capable of binding the CENP-B box motif (Supplementary Fig. 11). However, CENP-B boxes were not detected at satellite-free centromeres or within the TINSat1 and TINSat2 consensus sequences.

A genome-wide search for CENP-B boxes revealed that nearly all satellite-based centromeres either completely lack CENP-B boxes or contain only a few scattered sites, located at distances incompatible with a functional role in CENP-B recruitment (Supplementary Table 13). TIN25 showed slightly higher numbers, but the motifs were scattered within pericentromeric regions not bound by CENP-A and interspersed with telomeric-like repeats (Supplementary Table 13). A notable exception is TIN9, in which both haplotypes 1 and 2 contain 167 and 210 CENP-B boxes, respectively. These boxes are contained within a 230 kb region of TINSat1 arrays not bound by CENP-A (Fig. 6A, Supplementary Table 13). In haplotype 1, this array lies at the edge of the pericentromeric region, whereas in haplotype 2 it represents the sole satellite sequence within the misassembled centromeric locus. Immunofluorescence analysis using an anti-CENP-B antibody confirmed that the only labeled primary constriction corresponds to chromosome 9. Signal intensity varied between homologs, suggesting potential copy number differences (Fig. 6B). The CENP-B box containing arrays show low sequence identity with the CENP-A binding array of chromosome 9 (Fig. 6A) and of other chromosomes (Fig. 2B), suggesting that it may represent a distinct variant of the TINSat1 satellite. Tandem Repeat Finder analysis confirmed that they consist of a TINSat1 variant with 682 bp monomers, each carrying a CENP-B box (Fig. 6C, Supplementary Fig. 12). To avoid potential bias in estimating the genomic abundance of this variant due to the reference assembly, we aligned our ChIP-seq reads to both TINSat1 variant sequences and found that the CENP-B box-containing variant accounts for only 2.6% of all TINSat1 copies and is not enriched in CENP-A bound chromatin (Fig. 6D, Supplementary Table 14).

**Figure 6.**
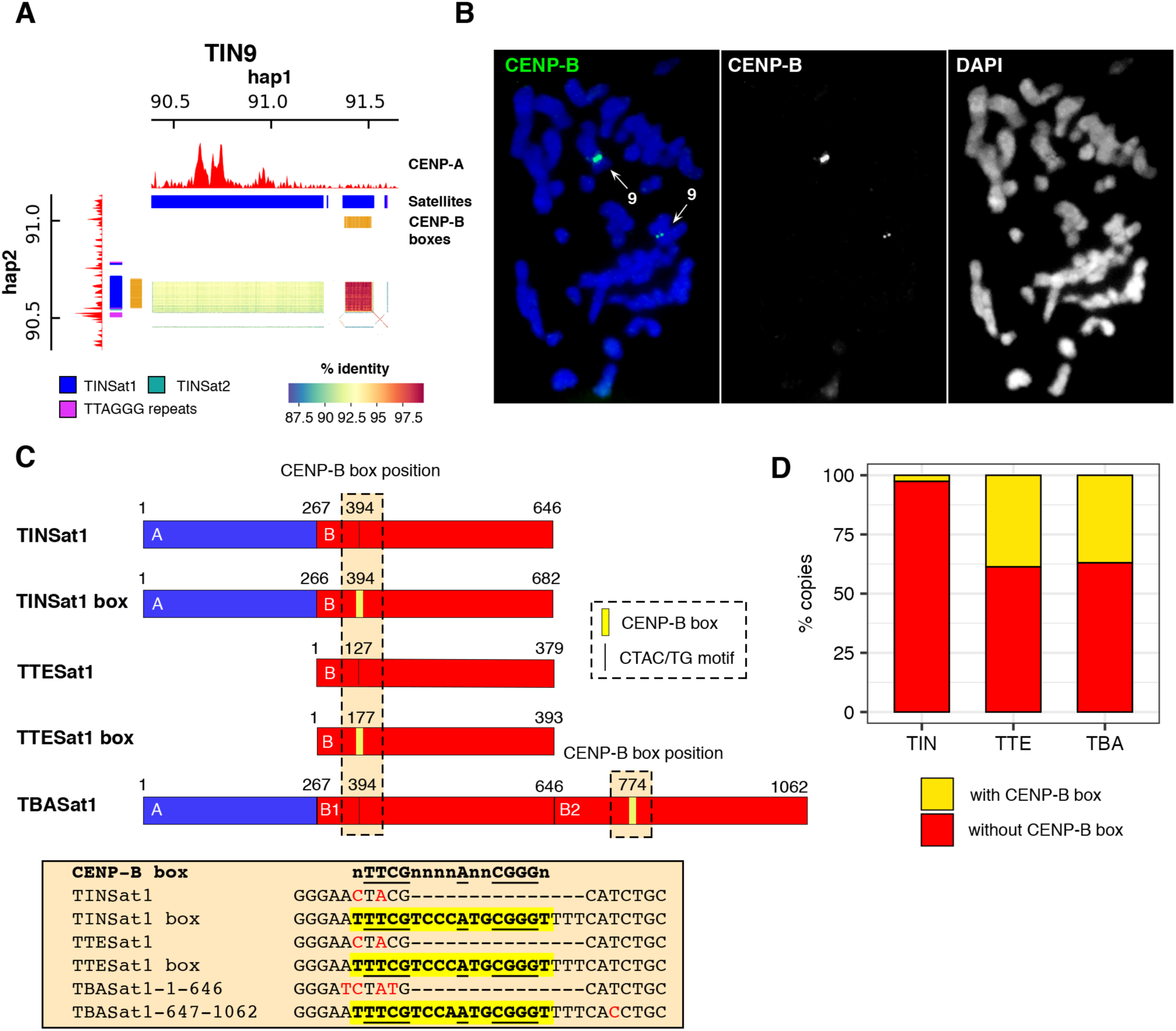
CENP-B binding in *Tapirus* species. A) Identification of CENP-B boxes within TINSat1 arrays on chromosome 9. ChIP-seq profiles of CENP-A in mTapiInd1.hap1 (hap1) and mTapiInd1.hap2 (hap2) are shown. The centromeric locus is misassembled in haplotype 2. Colored bars beneath the CENP-A peaks represent satellite DNA arrays, with colors corresponding to satellite families defined in the legend. Clusters of CENP-B boxes are shown in orange. Sequence identity maps between the two haplotypes, generated using ModDotPlot, are displayed in the center. B) Immunofluorescence with an anti-CENP-B antibody on *T. indicus* metaphase chromosomes. CENP-B signals (green) are detected on the pair of chromosome 9, indicated by arrows. C) Schematic representation of Sat1 satellite monomers identified in *T. indicus* (TINSat1), *T. terrestris* (TTESat1) and *T. bairdii* (TBASat1). Variants with and without the CENP-B box are shown, with regions of sequence identity indicated in different colors. The position of the CENP-B box - replaced by a CTAC/TG motif in the variants lacking it - is highlighted. Below the schematic, an alignment of the regions with and without the CENP-B box across the different monomers is shown. D) Abundance of the monomer variants containing (yellow) or lacking (red) the CENP-B box in the genomes of *T. indicus* (TIN), *T. terrestris* (TTE) and *T. bairdii* (TBA).

To investigate the evolutionary history of this satellite family in relation to the presence of the CENP-B box, we examined the satellite families identified by TAREAN in the other tapir species (Supplementary Table 8). Satellites homologous to TINSat1 and TINSat2 were found in both *T. terrestris* and *T. bairdii*, suggesting that the Sat1 and Sat2 satellite families were already present in the common ancestor of tapirs (Fig. 6D, Supplementary Fig. 12 and 13). Sat1 is the most abundant satellite family in both species (Supplementary Table 8). Despite the high sequence identity among Sat1 elements across tapir species (Supplementary Fig. 13), monomer length varies: in *T. terrestris*, the TTESat1 repeat unit is 379 bp, whereas in *T. bairdii*, the TBASat1 monomer is 1062 bp (Supplementary Table 8). Sequence analysis of the Sat1 units from the three species revealed the presence of distinct subregions (Fig. 6C). The TTESat1 monomer is found in all three species: once in *T. indicus* and in two tandem copies in *T. bairdii*, likely as a result of a duplication event. An additional region is present as a single copy only in *T. indicus* and *T. bairdii*.

The search for the CENP-B box motif showed that, in *T. terrestris*, the primary TTESat1 consensus sequence identified by TAREAN lacks a CENP-B box, while the second most abundant consensus variant includes it. In *T. bairdii*, the CENP-B box is present only in the second copy of the duplicated region. Alignment of the Sat1 units of the three species confirmed that the CENP-B box is present at the same position in the box-containing units while it is replaced by a CTAC/TG motif in the box-lacking units (Fig. 6C, Supplementary Fig. 12). Mapping genomic reads to box-containing and box-lacking consensus sequences showed that the CENP-B box variant is rare in *T. indicus*, but constitutes 38.7% and 37% of Sat1 copies in *T. terrestris* and *T. bairdii*, respectively (Fig. 6D).

## DISCUSSION

Thanks to the invaluable collections of the San Diego Frozen Zoo, we performed a molecular-level characterization of centromeres in the same *T. indicus* individual for which the Vertebrate Genome Project generated a haplotype-resolved, nearly gapless assembly. Because (peri)centromeric sequences vary substantially among individuals (Logsdon et al. 2024), analyzing the same animal with a diploid reference was essential to accurately delineate satellite loci and associated centromeres. We demonstrated that most centromeres are associated with TINSat1, the major satellite DNA family in this species. These centromeres are often embedded within large, megabase-sized satellite arrays, where the CENP-A binding region occupies only a limited portion. However, the length of the (peri)centromeric arrays varied considerably, both among different chromosomes and between homologs, confirming the high sequence heterogeneity.

A subset of satellite-based centromeres resides within arrays shorter than 100 kb, suggesting that these may represent early stages of satellite DNA expansion. An extreme case is observed at the centromere of TIN21, where in both homologs the CENP-A domain extends beyond the 22 kb TINSat1 array, encompassing non-satellite sequences and representing a centromere at the boundary between satellite-free and satellite-based configurations. A similar partially satellite-based centromere organization was recently described in the primate *Hoolock leuconedys* (Hartley et al. 2025), suggesting that such configuration may be more widespread among mammals. In *T. indicus*, the CENP-A domains of satellite-based centromeres typically localize to satellite repeats showing the highest sequence conservation relative to the surrounding pericentromeric array. According to current models of satellite DNA evolution, these conserved regions represent the most recently expanded repeats (Fry and Salser 1977). This arrangement was first described in primate centromeres (Alexandrov et al. 2001; Henikoff et al. 2015) and subsequently confirmed through recent centromere annotations in complete assemblies (Altemose et al. 2022; Logsdon et al. 2024). Differences between homologs in the position of CENP-A binding domains were observed in several *T. indicus* centromeres. Similar variation has also been reported for satellite-based centromeres in primates (Maloney et al. 2012; Mahlke et al. 2023; Logsdon et al. 2024), mouse (Arora et al. 2023) and horse (Cappelletti et al. 2019). However, such variation is not purely epigenetic, but also reflects underlying differences in the satellite DNA sequence itself.

Beyond the satellite-based centromeres, we also identified three satellite-free centromeres in *T. indicus*, representing the first report of such centromeres in a non-equid species within the order Perissodactyla. These centromeres are located in LTR-rich regions, consistent with the widespread abundance of transposable elements at centromeres in other animals - including equids - and in plants (Longo et al. 2009; Neumann et al. 2011; Tsukahara et al. 2012; Klein and O’Neill 2018; Nergadze et al. 2018; Yadav et al. 2018; Chang et al. 2019; Cappelletti et al. 2022; Hayashi et al. 2022; Hartley et al. 2025). The recurrent association of transposable elements with centromeres across diverse taxa may point to a potential role in centromere specification. Interestingly, the satellite-free centromere on TIN18 contains only very short stretches of satellite repeats, possibly representing an early stage of satellite DNA expansion. This pattern mirrors our previous observations in the X chromosomes of two zebra species, where satellite-free CENP-A binding domains include minimal satellite sequences, likely representing the initial seeds of satellite DNA acquisition (Cappelletti et al. 2022).

The first clear demonstration that the localization of CENP-A binding domains is not fixed, but that epialleles for CENP-A binding exist within populations and are inherited as Mendelian traits, came from our study on equid satellite-free centromeres (Purgato et al. 2015; Nergadze et al. 2018; Cappelletti et al. 2023). This phenomenon of centromere sliding is also present at the satellite-free centromeres of *T. indicus*. Notably, at the TIN15 centromere, the two epialleles are located 3.7 Mb apart on the two homologs, a positional shift significantly greater than the 600 kb sliding window observed in equids to date. Comparable megabase-scale shifts in centromere position have recently been reported in humans (Logsdon et al. 2024) and plants (Liu et al. 2023; Nagaki et al. 2025), indicating that centromeres can slide over broad genomic regions. The CENP-A enrichment profile of each *T. indicus* epiallele often displays a major peak separated from a minor peak by approximately 50 kb. This distinctive arrangement has also been observed at the satellite-free centromeres of certain horse individuals (Cappelletti et al. 2023), as well as at some satellite-based centromeres in humans (Altemose et al. 2022), and may reflect a specific organization of centromeric chromatin within nuclear architecture.

The satellite-free centromeres of *T. indicus* emerged following centromere repositioning events. Surprisingly, two of these centromeres are located in regions orthologous to centromeric regions of horse and donkey chromosomes that also carry satellite-free centromeres. In particular, the CENP-A binding domain of horse chromosome 11 and tapir chromosome 4 are precisely located at orthologous positions. In contrast, the centromere of donkey chromosome 8 has accumulated lineage-specific DNA duplications (Nergadze et al. 2018), which likely represent early stages of satellite DNA emergence. These duplications, which are bound by CENP-A in the donkey, occur within the region orthologous to the interval separating the TIN15 epialleles. Comparative analysis of orthologous chromosomes in other Perissodactyla species revealed that these satellite-free centromeres independently originated in the Malayan tapir, horse, and donkey as a result of distinct centromere repositioning events. Taken together, these findings suggest the existence of genomic hotspots for neocentromere formation within the Perissodactyla clade. Hotspots for neocentromere formation were identified in the human genome, where certain regions are recurrently involved in clinical neocentromere formation (Marshall et al. 2008). In addition, some clinical neocentromeres arise in genomic regions proximal to the sites where evolutionary neocentromeres associated with satellite DNA were seeded in other mammals, particularly primates (Ventura et al. 2003; Ventura et al. 2004; Cardone et al. 2006; Capozzi et al. 2008; Capozzi et al. 2009; Rocchi et al. 2012). These observations suggest that specific genomic locations are predisposed to centromere formation, either due to inherent sequence features or epigenetic context. The hotspots described in this study - particularly the TIN4/ECA11 region - represent striking examples where, despite more than 56 million years of divergence, the CENP-A binding domains are maintained at the same conserved genomic position across species.

Another point of similarity between *T. indicus* and equid species is the uncoupling between CENP-A and CENP-B, the only centromeric protein known to bind a specific DNA motif, whose functional role at centromeres remains elusive. In equids, CENP-B binding is limited to a few pericentromeres or to non-centromeric sites (Cappelletti et al. 2025b). Similarly, in *T. indicus*, CENP-B binds the pericentromeric region of only one chromosome pair. In this species, a variant of the major centromeric satellite lacking the CENP-B box is predominant over the box-containing variant. By contrast, in other tapir species, the two satellite variants - with and without the CENP-B box - are approximately equally represented in the genome. Interestingly, *T. indicus* is the only known species within Ceratomorpha to exhibit both extensive karyotypic plasticity over evolutionary time and remarkable centromere plasticity, as evidenced by the coexistence of satellite-based and satellite-free centromeres, along with non-centromeric satellite loci that may represent relics of inactivated ancestral centromeres. This observation supports our model linking evolutionary centromere plasticity to the absence of CENP-B binding at centromeres (Cappelletti et al. 2025b). Given that most studies on CENP-B have focused on humans and mice (Fachinetti et al. 2015; Kumon et al. 2021) - species in which all centromeres, except that of the Y chromosome, are bound by this protein - our findings underscore the importance of investigating this enigmatic protein in other mammals to unravel its role in the biodiverse landscape of centromere organization.

## METHODS

### Cell lines

The primary fibroblast cell line from a female *T. indicus* individual (GAN: MIG12-30034834) was provided by San Diego Zoo Wildlife Alliance’s Frozen Zoo. Fibroblasts were cultured in high-glucose DMEM medium, supplemented with 20% fetal bovine serum, 2 mM glutamine, 2% non-essential amino acids, and 1% penicillin/streptomycin. Cells were maintained in a humidified atmosphere of 5% CO2 at 37 °C.

### Identification and annotation of satellite repeats

Identification of satellite repeats from unassembled whole genome sequencing reads was performed with TAREAN (Galaxy Version 2.3.8.1), a computational pipeline that uses graph-based repeat clustering to detect satellite repeats directly from unassembled short reads (Novák et al. 2017) using 2 million reads as sample size and default parameters. For *T. indicus*, the analyzed reads were a subsample of the non immunoprecipitated input dataset. For the other Perissodactyla species, we used whole-genome sequencing datasets available in NCBI SRA Archive whose accession numbers are reported in Supplementary Table S8. Subsampling was carried out using SeqKit (2.6.1 version) (Shen et al. 2016). The identified satellite repeats were mapped on the used reference genomes using RepeatMasker (Galaxy Version 4.0.9) available at the Galaxy web platform (Community 2022). Satellite regions of each family located within 2000 bp were merged using Bedtools v2.30.0. Sequence identity among satellite families of the genus *Tapirus* was assessed with Clustal Omega (Madeira et al. 2024), and the resulting similarity values were visualized as heatmaps using R packages. Sequence identity heatmaps were generated with ModDotPlot v0.9.0 (Sweeten et al. 2024). Self-comparisons of each sequence were obtained using the --bin-freq mode, resulting in self-identity heatmaps. In addition, pairwise comparisons between different satellite sequences were performed without the --bin-freq option. The identity values from these pairwise plots were averaged to produce a summary similarity matrix, which was then visualized as a clustered heatmap using standard Python libraries.

### Identification and annotation of telomeric repeats

Telomeric repeats in the *T. indicus* diploid assembly were identified with a BLASTN 2.11.0 search for the (TTAGGG)₄ motif using the parameters -evalue 10 -word_size 11 -gapopen 5 -gapextend 2 -reward 2 -penalty -3 -outfmt 7. Hits within 250 bp of each other were merged into single loci using Bedtools v2.30.0.

### ChIP-seq

Chromatin from *T. indicus* primary fibroblasts was cross-linked with 1% formaldehyde, extracted, and sonicated to generate DNA fragments ranging from 200 to 800 bp. Chromatin immunoprecipitation was performed as previously described (Nergadze et al. 2018), using an anti-CENP-A serum developed in our laboratory (Cappelletti et al. 2019). Paired-end sequencing was carried out on a NovaSeq 6000 platform (IGA Technology Services, Udine, Italy). Sequencing quality was assessed with MultiQC v1.12.

ChIP-seq reads were aligned in paired-end mode to the diploid *T. indicus* assembly (mTapInd1.hap2: GCA_031878705.1 and mTapInd1.hap1: GCA_031878655.1) with Bowtie2 v2.4.2 (Langmead and Salzberg 2012), using stringent parameters (*–very-sensitive –no-mixed –no-discordant –maxins 800*) as previously applied for mapping CENP-A ChIP-seq data on haplotype-phased assemblies (Jin et al. 2025). The resulting BAM files were filtered with Samtools v1.15.1 (Li et al. 2009) to exclude unmapped reads, supplementary alignments, and non-primary alignments (-F 2308).

ChIP-seq datasets from horse and donkey were previously described (Nergadze et al. 2018; Cappelletti et al. 2019)and are available in the NCBI SRA Archive (SRR9855368, SRR9855369, SRR9855370, SRR9855371, SRR5515970, and SRR5516015). Reads were aligned in paired-end mode with Bowtie2 v2.4.2 (Langmead and Salzberg 2012) to the TB-T2T assembly (GCF_041296265.1) for *E. caballus* and to the EquAss-T2T_v2 assembly (GCF_041296235.1) for *E. asinus*, using default parameters.

Normalized enrichment peaks were obtained with the bamCompare tool from the deepTools suite (3.5.0 version) (Ramírez et al. 2016) using RPKM normalization in subtractive mode. Plots were obtained with pyGenomeTracks (3.6 version) (Lopez-Delisle et al. 2021).

### Haplotype-resolved mapping of CENP-A binding domains at satellite-free centromeres

To distinguish CENP-A binding between homologous chromosomes at satellite-free centromeres, we exploited single-nucleotide variants (SNVs) differentiating the two haplotypes (mTapInd1.hap1 and mTapInd1.hap2). SNVs were identified with the MUMmer3 pipeline (Kurtz et al. 2004): haplotypes were aligned with nucmer, alignments were filtered with delta-filter to retain reciprocal-best hits, and variants were extracted with show-snps. Only substitutions located at least 20 bp away from indels or other SNVs were retained.

ChIP-seq alignments previously generated against the diploid genome were then filtered with Samtools v1.15.1 (Li et al. 2009) to retain only reads covering SNV positions without mismatches (“NM:0”; “150M” CIGAR). Because this filtering resulted in highly specific but sparse coverage, enrichment profiles were generated with the bamCompare tool from the deepTools suite v3.5.0 (Ramírez et al. 2016) using CPM normalization in subtractive mode, a bin size of 10 bp, and the – exactScaling option, parameters optimized for these filtered BAM files. Plots were obtained with pyGenomeTracks v3.6 (Lopez-Delisle et al. 2021).

### DNA sequence analysis at satellite-free centromeres

The DNA sequence composition of satellite-free centromeric domains was assessed in terms of both interspersed repetitive elements and GC content. Interspersed repeats were annotated with RepeatMasker (Galaxy Version 4.0.9) using the RepBase library (release October 26, 2018). The same analysis was extended to the entire *T. indicus* diploid genome to obtain genome-wide values for both haplotypes. Statistical significance was evaluated with a two-tailed Wilcoxon one-sample test by comparing centromeric regions with genome-wide values.

### Comparative genomic analysis

Pairwise alignments between whole genomes or whole chromosomes were performed with Chromeister v1.5.a (Pérez-Wohlfeil et al. 2019) using default parameters. The reference assemblies used were: the mTapInd1.hap2 assembly (GCA_031878705.1) for *T. indicus*, the TB-T2T assembly (GCF_041296265.1) for *E. caballus*, the EquAss-T2T_v2 assembly (GCF_041296235.1) for *E. asinus*, the Equus_quagga_HiC assembly (available at https://www.dnazoo.org) for *E. burchelli*, the Equus_grevyi_HiC assembly (available at https://www.dnazoo.org) for *E. grevyi*, the mDicBic1.mat.cur assembly (GCA_020826845.1) for *D. bicornis*, the ASM2365373v1 assembly (GCA_023653735.1) for *C. simum*, the Tapirus_terrestris_HiC (GCA_028533255.1) for *T. terrestris*, and the Tapirella_bairdii_HiC assembly (GCA_028533255.1) for *T. bairdii*.

Fine-scale orthologies were identified using BLAT v.36 (Kent 2002). Sequence conservation between homologous haplotypes of *T. indicus* and between orthologous centromeric domains was assessed by alignments with minimap2 2.22-r1101 (Li 2018). Homologous haplotypes were aligned with -x asm5 (∼5% divergence), whereas orthologous domains were aligned with -x asm20 -k15 -w10 to increase seeding sensitivity at higher divergence (∼20%). Resulting PAF files were visualized with pafplot v0.1.0

### Immunofluorescence

Metaphase spreads for immunofluorescence experiments were prepared according to standard procedures (Piras et al. 2010). Immunofluorescence was performed as previously described (Cappelletti et al. 2025b). Briefly, metaphase preparations were fixed with ice-cold methanol for 4 min, permeabilized in 1× PBS with 0.05% Tween-20 for 15 min at room temperature, and incubated at 37 °C for 2 h with either our anti-horse CENP-A serum (Cappelletti et al. 2019) or the anti-CENP-B antibody sc-22788 (Santa Cruz Biotechnology Inc.), both diluted 1:100. The anti-CENP-B antibody was raised against amino acids 535–599 at the C-terminus of human CENP-B, which are conserved in the *T. indicus* protein (Supplementary Figure S10).

### FISH

Metaphase spreads for fluorescence *in situ* hybridization were prepared using the standard air-drying procedure. Whole-genomic DNA from *T. indicus* fibroblasts was extracted according to standard protocols. The telomeric probe is a mixture of 1-20 kb long synthetic (TTAGGG)n fragments that was previously prepared in our laboratory (Bertoni et al. 1994). Nick translation with Cy3-dUTP (Enzo Life Sciences) and hybridization were performed as previously described (Piras et al. 2010). Briefly, 100 ng of labelled whole-genomic DNA was used per slide. Hybridization of the genomic probe was performed under high-stringency conditions, while the telomeric probe was hybridized under low-stringency conditions as previously described (Piras et al. 2009; Piras et al. 2010).

Digital grayscale fluorescence images were acquired with a Zeiss Axio Scope.A1 fluorescence microscope equipped with a cooled CCD camera (Teledyne Photometrics). Pseudocoloring and image merging were performed using IPLab 3.5.5 Imaging Software. Chromosomes were identified by computer-generated reverse DAPI banding according to published karyotypes (Houck et al. 2000; Trifonov et al. 2008).

### Genomic distribution and abundance of satellite variants containing the CENP-B box

CENP-B boxes (nTTCGnnnnAnnCGGGn) were searched in the diploid *T. indicus* assembly using FIMO (MEME Suite v5.5.8) (Grant et al. 2011). Satellite loci containing clusters of CENP-B boxes were further characterized with Tandem Repeat Finder v.09 (Benson 1999). Comparative analyses of box-containing versus box-lacking satellite variants were performed with MultAlin (Corpet 1988). To estimate the genomic abundance of each satellite variant across tapir species, whole-genome sequencing reads were mapped in single-end mode with Bowtie2 (Langmead and Salzberg 2012) to 150-bp consensus fragments derived from TAREAN and Tandem Repeat Finder analyses, centered on either the CENP-B box or the alternative CTAC/TG motif, flanked by 50 N bases on each side to minimize edge-mapping artifacts. Normalized read counts (CPM) were calculated. In *T. indicus*, both input and CENP-A ChIP-seq datasets were mapped to these consensus variants to determine their relative enrichment in the immunoprecipitated fraction.

## ACKNOWLEDGEMENTS

This research was funded by Animal Breeding and Functional Annotation of Genomes (A1201) Grant 2019–67015-29340/Project Accession 1018854 from the USDA National Institute of Food and Agriculture. The Galaxy server that was used for some calculations is in part funded by Collaborative Research Centre 992 Medical Epigenetics (DFG grant SFB 992/1 2012) and German Federal Ministry of Education and Research (BMBF grants 031 A538A/A538C RBC, 031L0101B/031L0101C de.NBI-epi, 031L0106 de.STAIR (de.NBI)). Computational resources for RepeatExplorer analysis were provided by the ELIXIR-CZ project (LM2023055), part of the international ELIXIR infrastructure.

